# Doublecortin reinforces microtubules to promote growth cone advance in soft environments

**DOI:** 10.1101/2024.02.28.582626

**Authors:** Alessandro Dema, Rabab A. Charafeddine, Jeffrey van Haren, Shima Rahgozar, Giulia Viola, Kyle A. Jacobs, Matthew L. Kutys, Torsten Wittmann

## Abstract

Doublecortin (DCX) is a microtubule-associated protein critical for brain development. Although most highly expressed in the developing central nervous system, the molecular function of DCX in neuron morphogenesis remains unknown and controversial. We demonstrate that DCX function is intimately linked to its microtubule-binding activity. By using human induced pluripotent stem cell (hiPSC)- derived cortical i^3^Neurons genome engineered to express mEmerald-tagged DCX from the endogenous locus, we find that DCX-MT interactions become highly polarized very early during neuron morphogenesis. DCX becomes enriched only on straight microtubules in advancing growth cones with approximately 120 DCX molecules bound per micrometer of growth cone microtubule. At a similar saturation, microtubule-bound DCX molecules begin to impede lysosome transport, and thus can potentially control growth cone organelle entry. In addition, by comparing control, DCX-mEmerald and knockout DCX -/Y i^3^Neurons, we find that DCX stabilizes microtubules in the growth cone peripheral domain by reducing the microtubule catastrophe frequency and the depolymerization rate. DCX -/Y i^3^Neuron morphogenesis was inhibited in soft microenvironments that mimic the viscoelasticity of brain tissue and DCX -/Y neurites failed to grow toward brain-derived neurotrophic factor (BDNF) gradients. Together with high resolution traction force microscopy data, we propose a model in which DCX-decorated, rigid growth cone microtubules provide intracellular mechanical resistance to actomyosin generated contractile forces in soft physiological environments in which weak and transient adhesion-mediated forces in the growth cone periphery may be insufficient for productive growth cone advance. These data provide a new mechanistic understanding of how DCX mutations cause lissencephaly-spectrum brain malformations by impacting growth cone dynamics during neuron morphogenesis in physiological environments.

## Introduction

Doublecortin (DCX) is a microtubule (MT)-associated protein exclusively expressed in immature neurons and essential for early brain development^1^. Mutations in the DCX gene account for nearly a quarter of all cases of lissencephaly-spectrum neurodevelopmental disorders^2,3^, malformations of the brain cortex that result in cognitive and motor impairments as well as epilepsy. Because DCX is encoded on the X-chromosome, the clinical manifestations of DCX mutations vary broadly from nearly normal in females to agyria, the complete absence of cortical folds, in affected males^4^. The underlying cause of cortical malformations is generally thought to be a failure of neuronal migration during brain development^5^. However, the molecular mechanism by which DCX supports the migration of immature neurons through the developing cortex is not understood.

DCX interacts with MTs through two conserved DCX domains that are connected by a flexible linker and are both required for MT association in cells and in vitro^6,7^. The two DCX domains are not equivalent and are thought to play related, yet distinct roles in MT nucleation and stabilization. In vitro experiments and computational modelling suggest that the N-terminal DCX domain is primarily required for MT association. In vitro, DCX is also a strong MT nucleator^8,9^, and the C-terminal DCX domain is implicated in this MT nucleation activity but may also play a role in cooperative DCX self-association during MT binding^7,9,10^. MTs are hollow tubes that assemble from α/β tubulin dimers arranged head-to-tail along protofilaments that laterally interact to form the MT wall^11,12^. Ultrastructural analysis demonstrated that DCX domains bind to the vertex between four tubulin dimers^13^. This unique binding site between adjacent MT protofilaments changes conformation and longitudinally shortens during GTP hydrolysis and is shared by EB1 family proteins that associate with growing MT plus ends. Although in vitro very low concentrations of DCX can recognize growing MT ends^10^, we previously reported that this is not the case in cells. In contrast, DCX specifically recognizes the compacted GDP MT lattice and in cells is excluded from the GTP/GDP-Pi cap that is recognized by EB1^6^. This preference of DCX for GDP-like MT wall conformations with compacted inter-dimer spacing is further supported by its dissociation from MTs with high local curvature^6^. DCX also preferentially binds MTs with thirteen protofilaments, the predominant eukaryotic MT geometry^14^, and both in vitro and in cells protects MTs from depolymerization^6,8^. Lastly, in vitro, DCX can _influence_ how motor proteins interact with MTs potentially affecting intracellular transport and organization^15^.

Thus, DCX influences many diverse aspects of MT function and dynamics. Yet, it remains unclear which of these DCX properties are relevant in developing neurons. Here we show that disruption of MT-binding is central to DCX pathology, and by using genome-edited human induced pluripotent stem cells (hiPSCs) that reliably differentiate into cortical neurons^16^, demonstrate that at endogenous expression levels DCX is nearly exclusively bound to and stabilizes growth cone MTs. Our results support a model in which DCX predominantly functions to mechanically reinforce the growth cone cytoskeleton to allow productive growth cone advance in physiological soft environments.

## Results

### Association with straight microtubules is central to doublecortin (DCX) function

Although the vast majority of pathogenic DCX missense mutations occur in one of the two conserved DCX domains^2,7^, it has not been rigorously assessed how these mutations alter DCX interactions with MTs in cells or how DCX-MT binding relates to disease phenotypes. To quantitatively compare MT-binding of a range of clinically relevant, pathogenic DCX mutations, we selected missense mutations for which varying degrees of cortical malformation phenotypes have been documented^2,17–20^ and that were mostly distributed throughout the N-terminal DCX domain, primarily responsible for MT-binding^7^, as well as mutations in the first loop of the C-terminal DCX domain.

Quantitative comparison of EGFP-tagged MT-bound DCX with fluorescent signal in the surrounding cytoplasm in non-neuronal RPE cells expressing similar low levels of these DCX constructs (Fig. S1) showed that all sixteen missense mutations tested reduced DCX-MT binding highly significantly regardless of whether cryo-EM data predicted these mutations to directly participate in interactions with the MT wall or if they were located elsewhere^7^ (Fig. 1a, c). In addition, as far as this was possible to discern for DCX constructs with weak MT-binding such as N94D, which is located in a loop that defines the main structural difference between the two DCX domains, mutations with remaining MT-binding also remained specific for straight MTs in cells. This is important as it indicates that the specificity for straight MTs is not a characteristic of a single DCX domain by itself but a property of the DCX molecule containing both DCX domains that remains to be fully understood.

**Figure 1.**
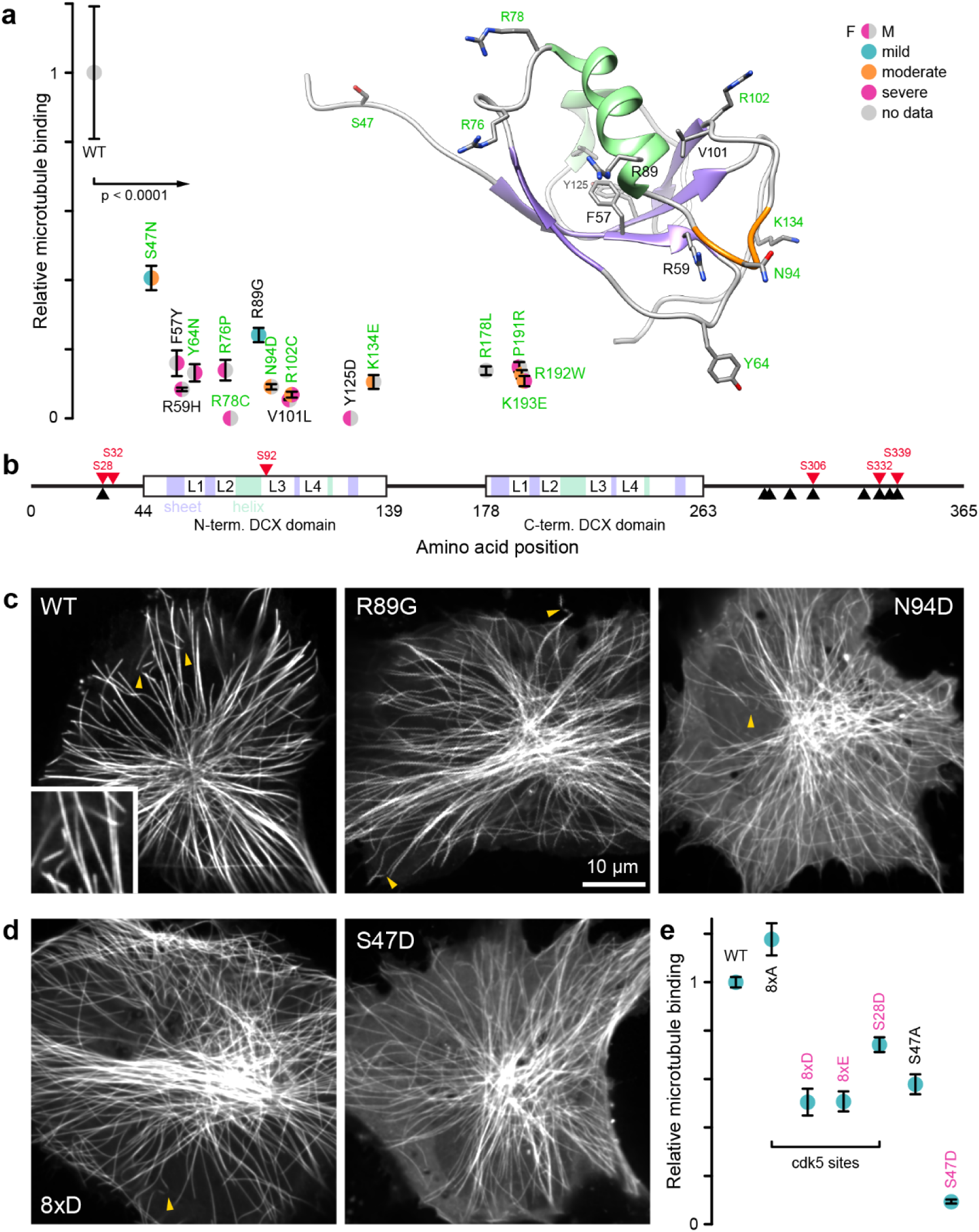
Pathogenic missense mutations disrupt DCX-binding to MTs. **(a)** Quantification of MT-binding of EGFP-tagged DCX with the indicated missense mutations in transiently transfected RPE cells. Data are from 3 independent experiments with error bars indicating standard deviation and 25-30 cells analyzed for each DCX construct. Relative MT-binding is the ratio of MT-bound to local cytoplasm fluorescence intensity compared to wild-type (WT) DCX-EGFP. Symbols are color-coded to reflect the clinical severity of specific mutations. Inset shows the structure of the N-terminal DCX domain viewed from the MT-binding surface. Residues labelled in green are predicted to directly interact with the MT wall. The loop shown in orange is unique to the N-terminal DCX domain. All mutations show highly significantly decreased MT-binding by ANOVA with Tukey-Kramer HSD. **(b)** DCX domain structure. Black triangles indicate the position of putative cdk5 phosphorylation sites, red triangles are phosphorylated residues identified by mass spectrometry in DCX-mEmerald i^3^Neurons. **(c)** Representative images of indicated DCX-EGFP constructs in RPE cells. Yellow arrows highlight MT segments with high local curvature and decreased DCX-MT association. **(d)** Representative images of the indicated phosphorylation site mutations. **(e)** Quantification of MT-binding of the indicated phosphorylation site mutations relative to wild-type DCX-EGFP.

Due to the variability of clinical classification criteria in human patients with DCX-related cortical malformations and the small numbers of clinical reports for specific DCX mutations, it was difficult to clearly correlate the amount of MT-binding with the severity of clinical manifestations. Most moderate to severe DCX mutations clustered around 10-20% remaining MT-binding activity compared with wild-type DCX (Fig. 1a). However, notably the two mutations that displayed no residual MT-binding (R78C and Y125D) were associated with the most severe clinical phenotype in females and no data in males possibly due to non-viability, while the mildest clinical phenotypes displayed the highest level of remaining MT binding (S47N and R89G). Together these results demonstrate that DCX-binding to straight MTs is a primary DCX function that is disrupted in DCX-mediated cortical malformations.

### DCX dynamics in developing human neurons

Because DCX is only expressed in the developing nervous system, to determine the dynamics of DCX-MT interactions in developing neurons, we integrated an mEmerald fluorescent protein tag at the C-terminus of the endogenous DCX locus by CRISPR/Cas9 genome editing in i^3^N cells (Fig. 2a), an hiPSC line that expresses Ngn2 under a doxycycline-induced promoter to allow inducible differentiation into cortical glutamatergic i^3^Neurons^21^. Because i^3^N cells are derived from a male donor, these cells only have one copy of the DCX gene. Both genomic PCR and immunoblotting demonstrated replacement of DCX with DCX-mEmerald in the edited i^3^N line (Fig. 2b, c), both proteins were expressed at similar levels, and as expected DCX protein amounts increased rapidly after Ngn2 induction in both control and edited i^3^Neurons. DCX-mEmerald i^3^Neurons developed normally, and the average neurite length was indistinguishable from control i^3^Neurons (Fig. 2e).

**Figure 2.**
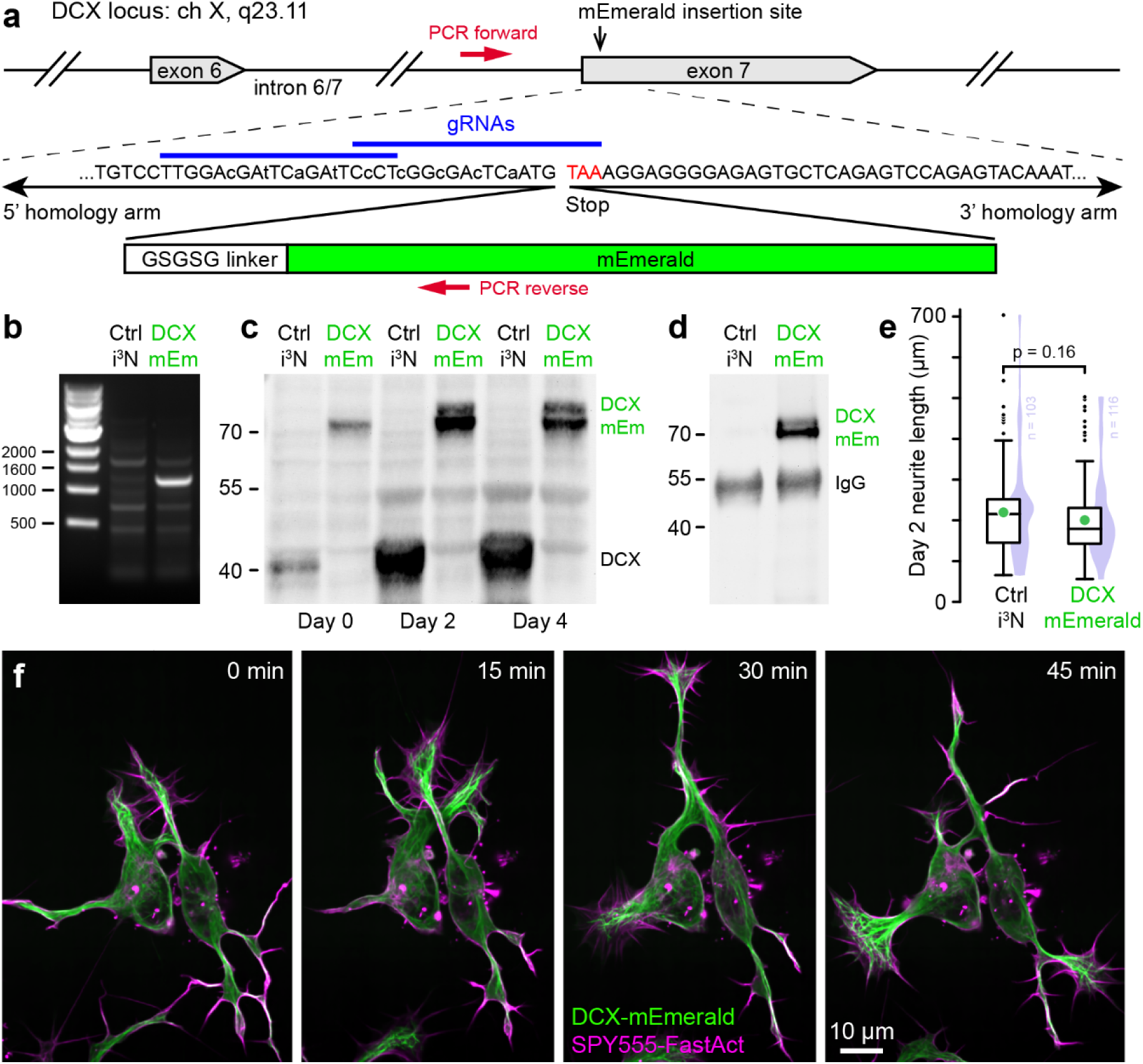
DCX-mEmerald expressed from the endogenous locus dynamically binds MTs in nascent growth cones. **(a)** Overview of the CRISPR/Cas9 genome editing strategy to insert mEmerald at the C-terminus of the endogenous DCX locus. Lowercase letters indicate silent sgRNA-resistant base pair changes in the 3’ homology arm. **(b)** Genomic PCR to validate mEmerald integration with primers as indicated in **(a)**. Only the edited i^3^N hiPSC line yields a PCR product of the expected size. **(c)** Immunoblot analysis of DCX expression in control and DCX-mEmerald edited i^3^Neurons at the indicated time points during i^3^Neuron differentiation showing the absence of untagged DCX in DCX-mEmerald i^3^Neurons and a similar increase of DCX protein in control and edited i^3^Neurons. **(d)** Immunoblot of an anti-GFP immunoprecipitation from control and DCX-mEmerald i^3^Neurons showing the expected ∼70 kDa DCX-mEmerald protein only in the edited line. **(e)** Comparison of the longest neurite lengths per cell in control and DCX-mEmerald i^3^Neurons two days after inducing differentiation. Box plots include the mean (green circle), and violin plots show the distribution of all measurements. Statistical analysis by unpaired t-test. **(f)** Time-lapse sequence of DCX-mEmerald i^3^Neurons labelled with SPY555-FastAct in very early stages of neurite outgrowth showing polarized DCX enrichment on MTs in nascent growth cones.

Remarkably, DCX-mEmerald localization to MTs became highly polarized very early during i^3^Neuron morphogenesis. Within hours of plating doxycycline-induced DCX-mEmerald i^3^N cells on laminin-coated plates, MTs invading nascent F-actin rich protrusions were brightly decorated with DCX-mEmerald, and DCX-MT binding disappeared when these temporary growth cones retracted (Fig. 2f; Video 1). DCX-mEmerald was similarly enriched on sometimes individual MTs invading collateral neurite branches (Video 2). This polarity persisted and at later stages of neuron development, DCX-mEmerald predominantly localized to growth cone MTs and was largely absent from MTs in the cell body or the neurite shaft (Fig. 3b). Similar to what we observed in non-neuronal cells, DCX-mEmerald predominantly bound straight growth cone MTs and reversibly dissociated from regions of high MT curvature (Fig. 3a; Video 3). Paclitaxel reversed this preference for straight MTs (Fig. 3c) and DCX-mEmerald was absent from the EB1-labelled domain at growing MT ends (Fig. 3d), consistent with our previous conclusions that DCX specifically recognizes straight GDP-MTs^6^.

**Figure 3.**
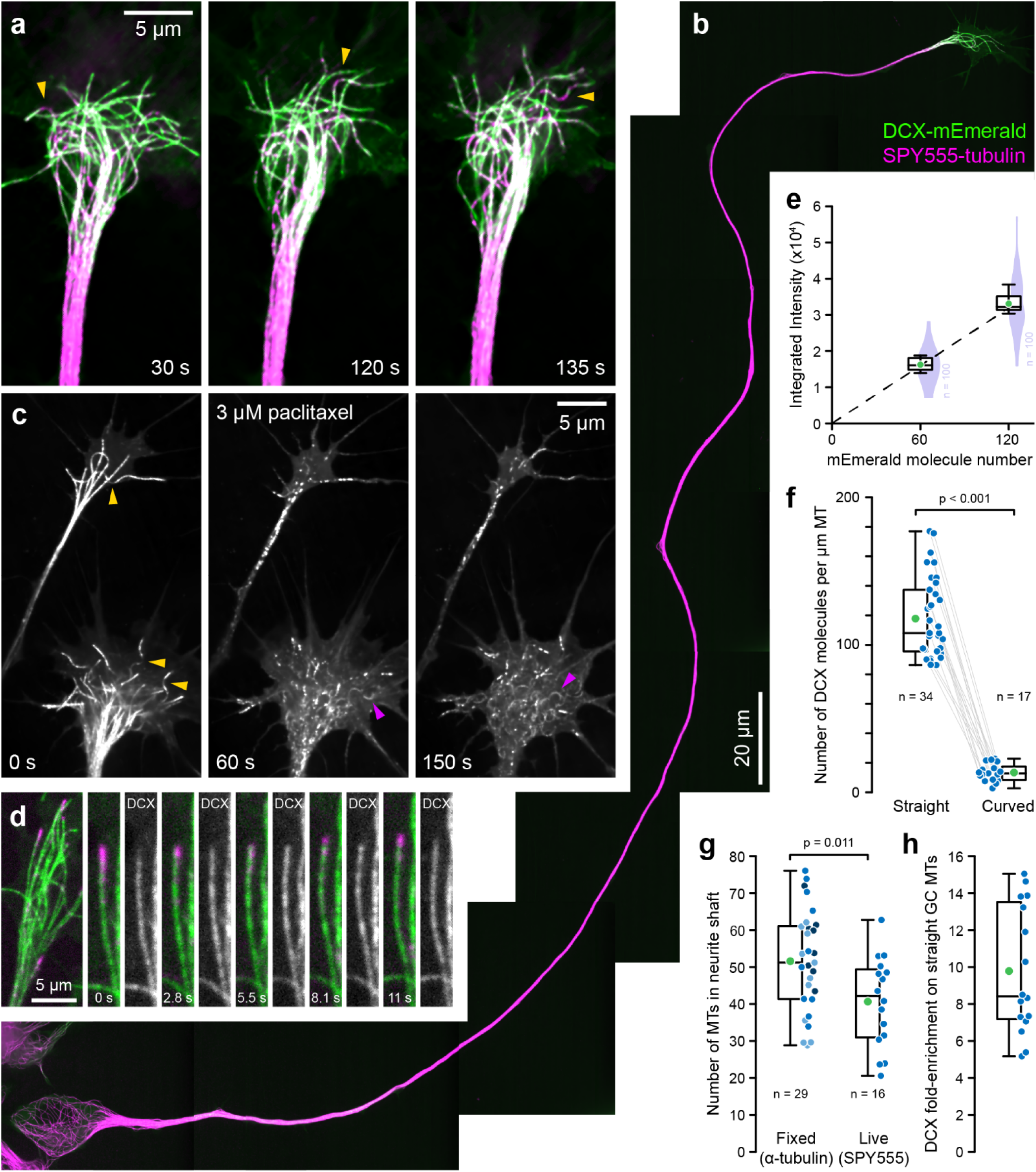
DCX is highly enriched on straight growth cone MTs. **(a)** Time-lapse sequence of a DCX-mEmerald i^3^Neuron growth cone labelled with SPY555-tubulin. Yellow arrowheads highlight MT buckling and dissociation of DCX-mEmerald from MTs with high local curvature. **(b)** DCX-mEmerald i^3^Neuron with a single long axon labelled with SPY555-tubulin illustrating exclusive DCX enrichment on growth cone MTs. Shown is a composite of eleven individual images. **(c)** Time-lapse of DCX-mEmerald i^3^Neuron growth cones after taxol addition showing rapid DCX dissociation from straight MTs and relocalization to high curvature MT segments (purple arrowheads). **(d)** Growth cone MTs in a DCX-mEmerald i^3^Neuron expressing EB3-tagRFP illustrating absence of DCX form the GPT/GDP-Pi cap at growing MT ends. **(e)** Calibration curve relating the number of mEmerald molecules on I3 nanocages to fluorescence intensity. The dashed line is a linear regression through the medians of 100 nanocages measured in 10 cells for each of the nanocage constructs and the origin (R^2^ = 1) used to calculate the number of DCX-mEmerald molecules from fluorescence intensity along growth cone MTs. Violin plots show the distribution of all nanocage intensity measurements. **(f)** Quantification of the number of DCX-mEmerald molecules on straight and curved growth cone MT segments. Grey lines connect measurements from the same growth cones. **(g)** Quantification of the number of MTs per neurite shaft in fixed (different shades indicate independent experiments) or live DCX-mEmerald i^3^Neurons by comparing the fluorescence intensity of a tubulin label on individual growth cone MTs with fluorescence intensity in the neurite shaft. **(h)** Calculation how enriched DCX-mEmerald is on straight growth cone MTs compared with MTs in the neurite shaft using the same SPY555-tubulin labeled live DCX-mEmerald i^3^Neuron dataset as in **(g)**. In **(f-h)**, box plots include the mean (green) and all individual data points (blue). Statistical analysis by unpaired t-test.

### DCX is highly enriched on growth cone microtubules

Because in the edited i^3^Neurons DCX-mEmerald is expressed from the endogenous promoter and therefore present at physiological protein levels, we measured how many DCX molecules are bound to growth cone MTs. By comparing the integrated DCX-mEmerald fluorescence intensity on growth cone MT segments with a fluorescence calibration curve generated by measuring the fluorescence intensity of two different I3 dodecahedron nanocage constructs with known numbers of mEmerald molecules^22,23^ (Fig. 3e; Fig. S2), we calculated that straight growth cone MTs bound 118 +/- 27 DCX-mEmerald molecules per µm MT (Fig. 3f). In contrast, the number of DCX-mEmerald molecules on curved MT segments was at least 10-fold reduced.

To further determine how enriched DCX is on growth cone MTs compared with MTs in the neurite shaft, we estimated the number of neurite MTs by calculating the fluorescence intensity ratio of the neurite MT bundle to individual MTs in the growth cone. Because neurites vary in thickness some variability in MT number is expected. Nevertheless, two different labeling methods yielded close estimates of an average of 40-50 MTs in a developing i^3^Neuron neurite MT bundle, either in fixed i^3^Neurons by α-tubulin immunofluorescence (52 +/- 14 MTs) or using SPY555-tubulin in live i^3^Neurons (41 +/- 12) (Fig. 3g). By then directly comparing the growth cone MT to neurite DCX-mEmerald fluorescence intensity ratio in SPY555-tubulin-labeled i^3^Neurons, we found that DCX-mEmerald is 10 +/- 3-fold enriched on growth cone MTs (Fig. 3h) resulting in a similarly low number of DCX molecules per neurite MT as on curved MT segments in growth cones. Of note, we likely overestimate remaining DCX-binding to neurite MTs because, in contrast to individual growth cone MTs, we cannot distinguish MT-bound from cytoplasm fluorescence in the neurite.

DCX is phosphorylated by several different protein kinases in vitro or in overexpression experiments and phosphorylation frequently inhibits MT-binding of MT-associated proteins^24^. To test if this gradient in DCX MT-binding activity could be due to phosphorylation, we first determined which residues were phosphorylated under physiological conditions by mass spectrometry of DCX-mEmerald immunoprecipitated with an anti-EGFP antibody from developing i^3^Neurons (Fig. 2d). Although we were unable to obtain complete coverage of the DCX-mEmerald protein sequence, the most highly phosphorylated residues were previously identified cdk5 sites^25^ (S28, S306, S332, and S339). One of these cdk5 sites (S28) is near the N-terminus while the other eight reside in the unstructured C-terminal part of DCX (Fig. 1b), but their influence on DCX MT-binding in an intracellular context remains poorly understood. However, DCX-EGFP constructs in which either S28 or all eight C-terminal cdk5 sites were mutated into phosphomimetic aspartate or glutamate residues displayed only moderately reduced MT-binding in transfected RPE cells (Fig. 1d, e) that cannot explain the 10-fold difference we observed between growth cone and neurite MTs in i^3^Neurons. Although our mass spectrometry results did not cover S47, S47 is directly involved in MT-binding^7^, mutated in neurodevelopmental disease (Fig. 1a) and S47 phosphorylation was previously proposed to reduce DCX MT-binding in vitro^26^. We therefore also tested a phosphomimetic S47D DCX-EGFP single amino acid change, which reduced MT-binding to a much greater extent (9 +/- 1% of WT; Fig. 1e) than mutation of all the C-terminal cdk5 sites together. Thus, while phosphorylation of S47, which is not a target of cdk5, sufficiently reduces DCX-MT binding to explain the MT-association gradient in neurons, this also suggests that cdk5 is not a major regulator of DCX MT-binding in developing neurons.

### DCX-binding to MTs impedes peripheral lysosome distribution

Recent in vitro data indicate that DCX binding to purified MTs greatly inhibits both binding and MT plus end-directed movement of kinesin-1 (KIF5) motors^15^. Because kinesin-1 is the principal motor protein driving fast anterograde lysosome transport in neurons^27,28^, we asked if lysosome dynamics differ in the growth cone in which DCX-binding to MTs is enriched from lysosome transport along the neurite shaft. Lysosomes were labelled with SiR-lysosome, a cell-permeable far-red-tagged pepstatin A that binds cathepsin D and that we previously validated to be highly lysosome specific^29^. Of note, most large and brightly labelled lysosomes remained localized to i^3^Neuron cell bodies. Yet, smaller dynamic cathepsin D-positive lysosomes moved rapidly along developing neurites at a velocity of 1.4 +/- 0.3 µm s^-1^ (Fig. 4a, b; Video 4). In contrast, once lysosomes entered growth cones, they became much more stationary, and their average velocity was substantially reduced (0.35 +/- 0.12 µm s^-1^).

**Figure 4.**
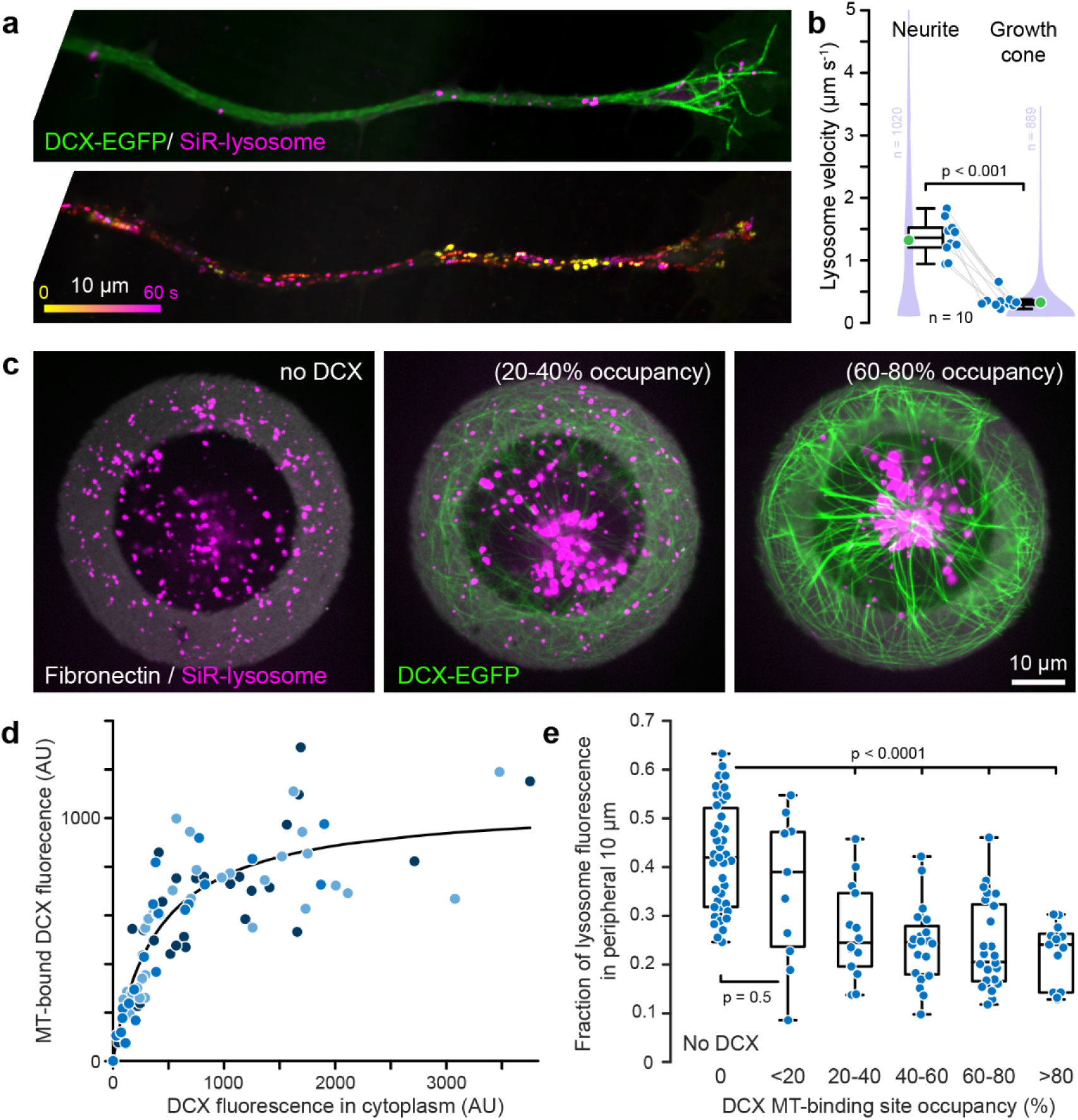
DCX can inhibit lysosome transport. **(a)** Lysosomes labelled with SiR-lysosome, a fluorogenic pepstatin derivative, in a DCX-mEmerald i^3^Neuron neurite. The bottom panel shows a color-coded projection of lysosome movements over 60 seconds. **(b)** Comparison of lysosome velocity in the neurite and the growth cone of DCX-mEmerald i^3^Neurons. Box plots include the mean (green) and all individual data points (blue). Grey lines connect data from the same i^3^Neurons. Violin plots show the distribution of all frame-to-frame lysosome velocity measurements. Statistical analysis by paired t-test. **(c)** Example images of RPE cells plated on 50 µm diameter, 10 µm wide fibronectin rings expressing no, low, or medium levels of DCX-EGFP. **(d)** DCX-MT-binding plotted as function of DCX-EGFP fluorescence signal in the cytoplasm. Each symbol is one cell with different shades indicating cells from three independent experiments (representing a total of 126 cells). The solid line is a fit with a hyperbolic binding curve indicating saturation of available MT binding sites. **(e)** Fraction of SiR-lysosome fluorescence in the peripheral 10 µm ring as a function of DCX-EGFP expression level binned into groups of similar DCX MT-binding site occupancy. Statistical analysis by ANOVA with Tukey-Kramer HSD.

Because the number of motile lysosomes in developing neurites was generally small and highly variable and this experiment also did not allow us to directly correlate the difference in lysosome dynamics in neurites and growth cones to the amount of DCX bound to MTs, we instead analyzed the distribution of lysosomes as a function of DCX-EGFP expression in transiently transfected RPE cells plated onto microprinted fibronectin rings (Fig. 4c)^30,31^. This standardizes cell shape and enables quantitative comparison of SiR-lysosome fluorescence distribution in many cells. By quantifying the amount of DCX-EGFP bound to MTs at different expression levels and fitting a saturation binding curve (Fig. 4d), we further estimated the DCX-EGFP expression level at which DCX-binding to MTs becomes saturated. This analysis showed that DCX-EGFP expression levels occupying as little as 20% of the available MT binding sites highly significantly inhibited lysosome distribution to the cell periphery (Fig. 4e) demonstrating that DCX can modulate intracellular transport.

### DCX is required for directed growth cone advance

Because of the clinical phenotype of DCX mutations, DCX is thought to be involved in the migration of immature neurons during brain development. However, the precise molecular function of DCX in developing neurons remains unknown. To test how DCX supports neuron development, we next generated DCX knockout i^3^Neurons by introducing a frameshift in exon 1 by CRISPR/Cas9 genome editing. As expected, these DCX -/Y cells no longer express DCX, but protein levels of MAPT/tau increased normally during i^3^Neuron differentiation indicating a normal transcriptional program (Fig. 5a). Given that DCX is dynamically enriched on MTs in nascent growth cones very early during i^3^Neuron morphogenesis (Fig. 2f, Videos 1 and 2), we expected a pronounced neurite outgrowth defect in DCX -/Y i^3^Neurons. To our surprise, however, DCX -/Y i^3^Neurons formed dynamic advancing growth cones similar to control and DCX-mEmerald i^3^Neurons (Fig. 5c), and on rigid tissue culture plastic, there was no obvious difference in morphology or neurite length between control, DCX-mEmerald and DCX -/Y i^3^Neurons. However, when we plated developing i^3^Neurons on soft 400 Pa polyacrylamide (PAA) gels that more closely resemble the tissue stiffness through which neurites elongate during brain development^32^, DCX -/Y neurites remained significantly shorter compared with control or DCX-mEmerald i^3^Neurons (Fig. 5b).

**Figure 5.**
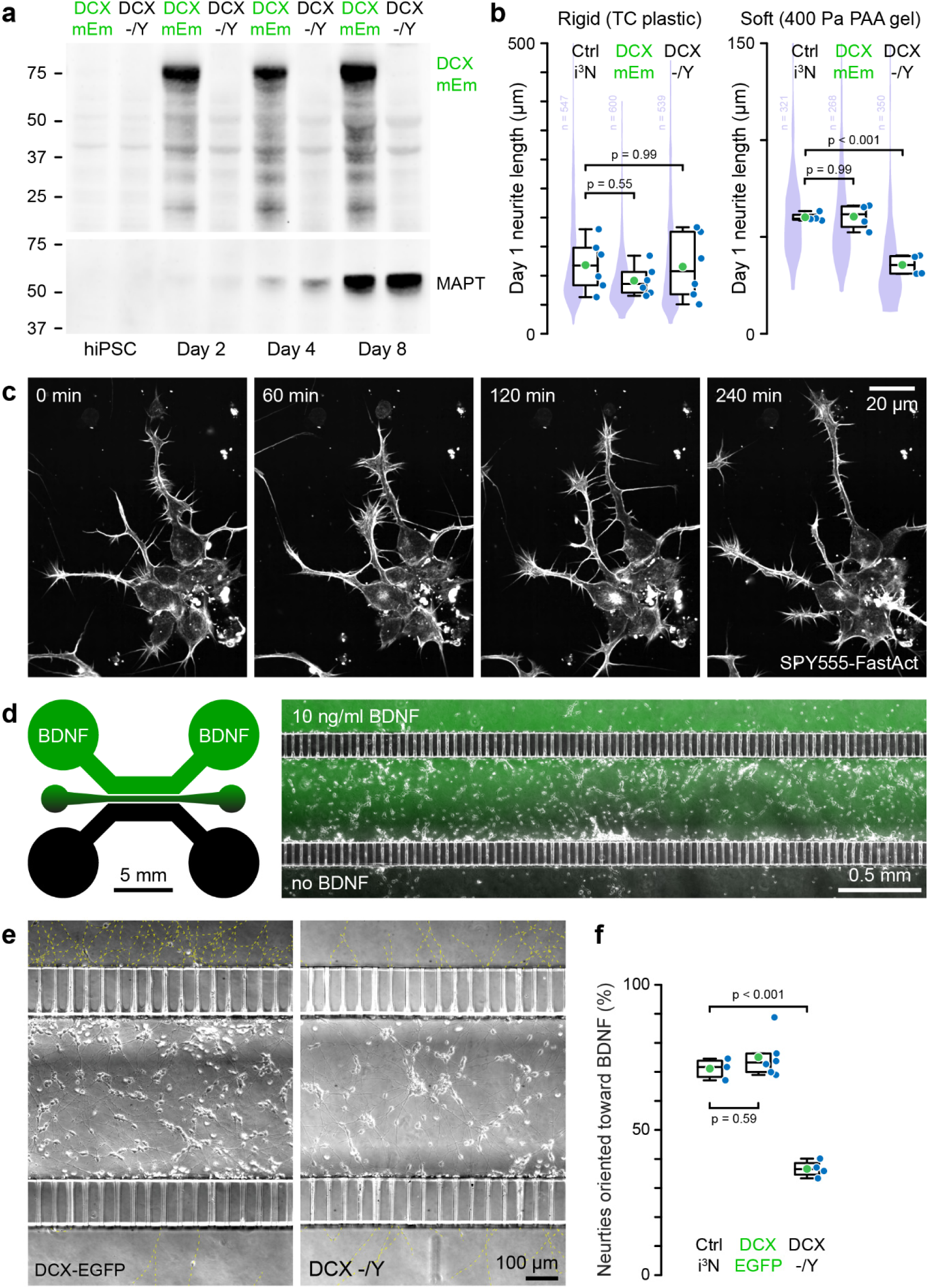
DCX is required for growth cone advance in physiological conditions. **(a)** Immunoblot analysis of DCX-mEmerald (top) and MAPT/tau (bottom) expression during i^3^Neuron differentiation in DCX-mEmerald and knockout DCX -/Y i^3^Neurons. **(b)** Comparison of neurite lengths of control, DCX-mEmerald and DCX -/Y i^3^Neurons on either rigid or soft substrates. Box plots include the mean (green) and all individual data points (blue) corresponding to the averages of measurements from six (rigid) and four (soft) independent experiments, respectively. Violin plots show the distribution of all neurite lengths measurements. Statistical analysis by ANOVA with Tukey-Kramer HSD. **(c)** Time-lapse sequence of SPY555-FastAct labelled DCX -/Y i^3^Neuron neurite outgrowth dynamics. **(d)** Diagram of the passive microfluidic device used to measure the chemotactic response of i^3^Neuron neurite growth to a BDNF gradient generated by diffusion across a central cell plating channel to a source channel with BDNF and a sink channel containing media without BDNF. The three channels are connected by microchannels (10 µm wide x 3 µm tall) that allow diffusion and neurites to grow through but confine cell bodies to the central channel. The representative image on the right shows control i^3^Neurons in the central channel and the BDNF gradient visualized with a fluorescent dextran of similar molecular weight as BDNF approximately 24 hours after plating. **(e)** Representative images of DCX-mEmerald and DCX -/Y i^3^Neurons in the central chemotaxis channel approximately 48 hours after plating. Neurites growing through the microchannels are highlighted by yellow dashed lines. The top source channel contains BDNF. **(f)** Quantification of the fraction of neurites growing into the BDNF source channel versus the sink channel after 24-48 hours. Box plots include the mean (green) and all individual data points (blue). Statistical analysis by ANOVA with Tukey-Kramer HSD.

Because dynamic MTs are required for growth cone guidance downstream of chemotactic signaling^33^, we next tested if in addition to this growth cone advance defect in physiological stiffness, DCX -/Y i^3^Neurons also display a growth cone guidance phenotype. Because growth cones sense gradients of brain-derived neurotrophic factor (BDNF)^34^, we explored growth cone chemotaxis in passive microfluidic chambers that allow formation of a BDNF gradient across a central cell plating channel (Fig. 5d). This central channel communicates with source and sink channels through microchannels that allow diffusion and neurites to grow through, but for the most part restrict movement of i^3^Neuron cell bodies. Counting the number of neurites that extended through the microchannels into either the source or sink channels revealed a clear bias of control and DCX-mEmerald i^3^Neuron neurites toward BDNF that was abolished in the DCX -/Y i^3^Neurons (Fig. 5e, f). Together, these data indicate that although a DCX -/Y phenotype was not immediately obvious in standard in vitro i^3^Neuron differentiation conditions, DCX is required for neurite elongation and growth cone guidance in more physiological environments.

### DCX protects growth cone MTs from depolymerization

Especially so-called ‘pioneer’ MTs that extend into the growth cone periphery are thought to be important for growth cone guidance^35^ and we next asked how the absence of DCX changes growth cone MT organization and dynamics. Indeed, pioneer MTs were shorter in DCX -/Y i^3^Neurons and, analyzed by measuring the distance of MT ends from the growth cone edge in fixed cells, did not penetrate as far into the growth cone periphery as in control and DCX-mEmerald i^3^Neurons (Fig. 6a, b).

**Figure 6.**
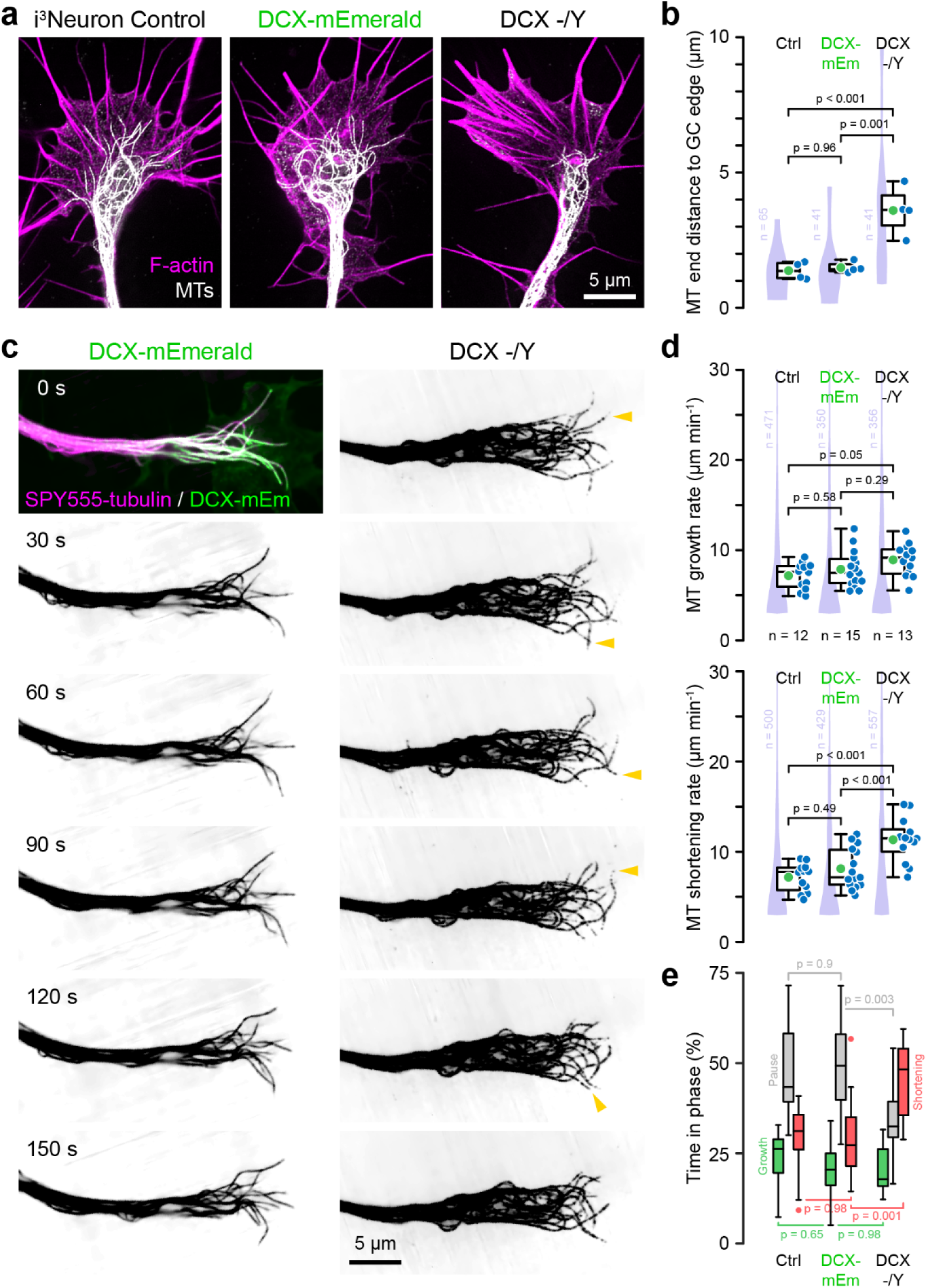
DCX stabilizes growth cone MTs. **(a)** Representative immunofluorescence images of MTs in control, DCX-mEmerald and DCX -/Y i^3^Neurons. **(b)** Comparison of the average distance between the ends of the three longest MTs from the growth cone edge. Box plots include the mean (green) and all individual data points (blue) corresponding to 10-15 growth cones from four independent experiments. Violin plots show the distribution of measurements from all growth cones. Statistical analysis by ANOVA with Tukey-Kramer HSD. **(c)** Time-lapse sequence of SPY555-tubulin labelled growth cone MT dynamics in DCX-mEmerald and DCX -/Y i^3^Neurons. To better visualize individual growth cone MTs, images are shown with inverted contrast and adjusted gamma. Yellow arrowheads indicate MTs in DCX -/Y i^3^Neurons that shorten frequently all the way back into the growth cone center, which is rarely seen in control or DCX-mEmerald i^3^Neurons. **(d)** Quantification of growth cone MT growth and shortening rates in control, DCX-mEmerald and DCX -/Y i^3^Neurons. Box plots include the mean (green) and all individual data points (blue) corresponding to the averages from individual growth cones. Violin plots show the distribution of all measurements. Statistical analysis by ANOVA with Tukey-Kramer HSD. **(e)** Quantification of the fraction of time growth cone MT ends spend in a growth, pause, or shortening phase with a pause defined as less than 150 nm (3 pixel) frame-to-frame displacement showing a shift toward more frequent shortening events in DCX -/Y i^3^Neurons. Statistical analysis by ANOVA with Tukey-Kramer HSD.

We next used a fluorogenic taxane derivative at very low concentrations to directly observe growth cone MT dynamics^16^ (Fig. 6c; Video 5). Although the MT growth rate appeared slightly elevated in DCX -/Y i^3^Neurons, this was not statistically significant. In contrast, the MT shortening rate was significantly increased in DCX -/Y (11.3 +/- 2.3 µm min^-1^) compared with control (7.2 +/- 1.5 µm min^-1^) and DCX-mEmerald i^3^Neurons (8.1 +/- 2.2 µm min^-1^) (Fig. 6d). In addition, and consistent with DCX inhibiting growth cone MT depolymerization, the fraction of time MTs spent in a shortening phase was nearly doubled in DCX -/Y i^3^Neurons while the time MTs spent growing was unaffected (Fig. 6e). Together these data indicate that DCX promotes and stabilizes pioneer MT extension into the growth cone periphery by protecting MTs from shortening events and slowing the depolymerization rate.

### DCX-stabilized MTs counteract growth cone compressive forces and enable productive traction

Because MTs bear compressive loads in cells^36^ and buckle due to coupling to F-actin retrograde flow^37,38^, which is similar to the buckling of DCX-covered MTs we observe in DCX-mEmerald i^3^Neurons (Fig. 3a; Video 3), we next asked how the reduction of growth cone MTs in DCX -/Y i^3^Neurons impacted growth cone F-actin dynamics. Using a fluorogenic jasplakinolide derivative, SPY650- FastAct, we measured F-actin retrograde flow rates (Fig. 7a, b), which in control (2.9 +/- 0.1 µm min^-1^) and DCX-mEmerald i^3^Neurons (2.9 +/- 0.04 µm min^-1^) were nearly identical to what we reported previously^16^. In contrast, F-actin retrograde flow was doubled in DCX -/Y i^3^Neurons (6.6 +/- 0.5 µm min^-1^) indicating that DCX-stabilized growth cone MTs counterbalance F-actin retrograde flow.

**Figure 7.**
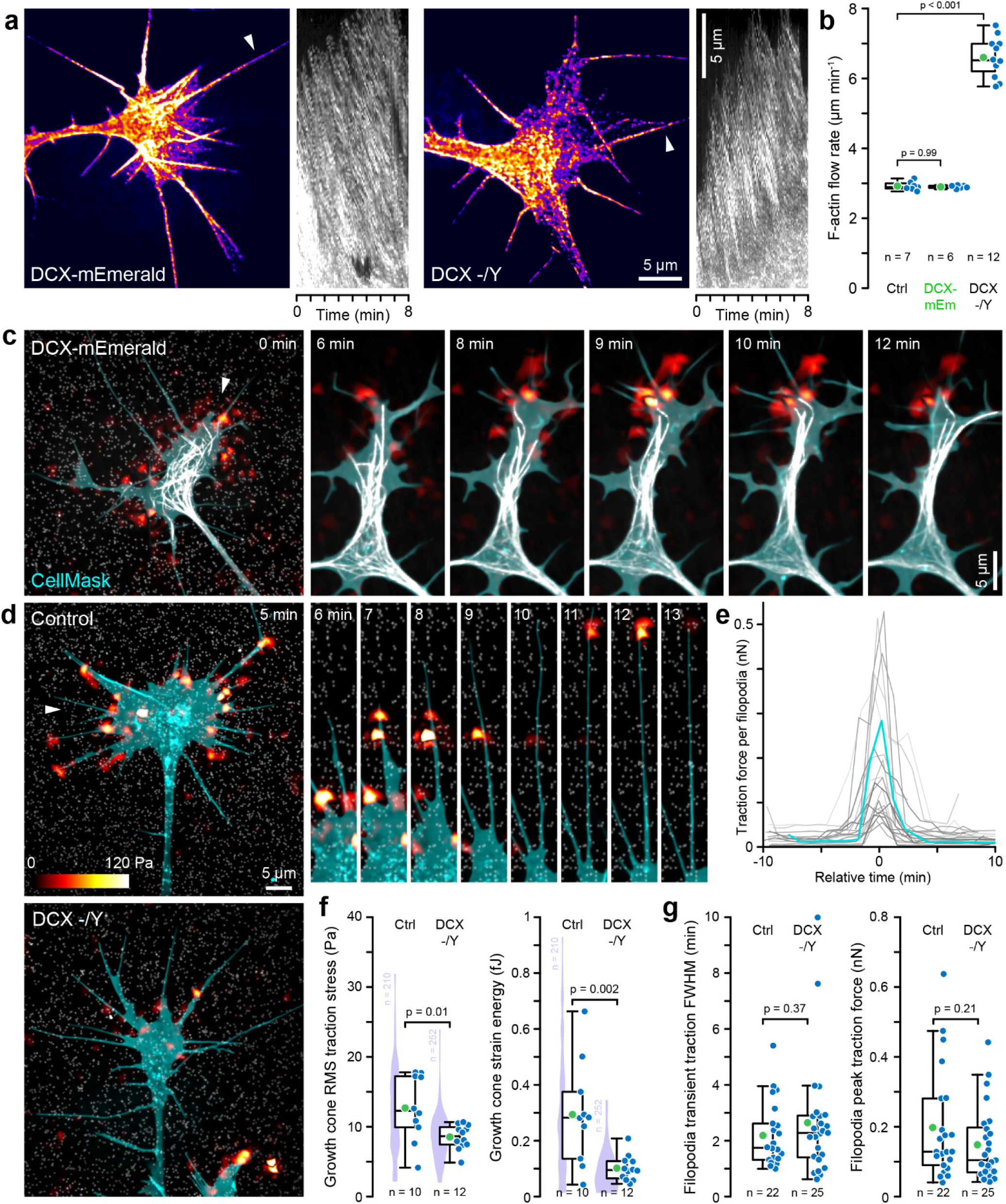
DCX stabilizes growth cone F-actin dynamics and traction forces. **(a)** Growth cones of DCX-mEmerald and DCX -/Y i^3^Neurons labelled with SPY650-FastAct and kymographs along the filopodia indicated by arrowheads. **(b)** Quantification of the F-actin retrograde flow rate in growth cones with and without DCX. Box plots include the mean (green) and all individual data points (blue) corresponding to at least three flow measurements per growth cone. Statistical analysis by ANOVA with Tukey-Kramer HSD. **(c)** Traction force microscopy of a representative DCX-mEmerald i^3^Neuron growth cone. DCX-mEmerald on growth cone MTs is white. For better visualization of these MTs, the polyacrylamide-embedded bead channel is omitted in the time-lapse sequence on the right. **(d)** Traction force microscopy of control and DCX -/Y i^3^Neuron growth cones. The time-lapse sequence shows transient traction stress peaks along the filopodia in the control i^3^Neuron. In both **(c)** and **(d)**, the CellMask-labelled growth cone is cyan, the polyacrylamide-embedded beads are shown in grey, and the traction stress map is shown with the same 0-120 Pa pseudo-color scale. The time-lapse sequences are 1.5x magnified and their location indicated by arrowheads in the images to the left. **(e)** Overlay of 22 filopodia traction force transients aligned to the peak force position determined by Gaussian fits. The cyan trace corresponds to the filopodium shown in the time-lapse in **(c)** at 7-9 min. **(f)** Quantification of whole growth cone root mean square (RMS) traction stress averages and strain energies. Box plots include the mean (green) and all individual data points (blue). Each data point is the average of 21 time points for each growth cone analyzed. Violin plots show the distribution of measurements from all time points of all growth cones. **(g)** Quantification of the full width half maximum (FWHM, i.e. duration) and peak forces of filopodia traction stress transients determined by Gaussian fits of individual force traces. Statistical analysis in **(f)** and **(g)** by unpaired t-test.

In addition to this intracellular linkage of F-actin to MTs^39^, F-actin retrograde flow is also inversely correlated to forward movement and traction forces exerted by cells through the adhesion clutch^40^. To better understand how stabilization of growth cone MTs by DCX contributes to force generation during growth cone advance, we performed time-lapse traction force microscopy (TFM) on 400 Pa polyacrylamide gels with embedded sub-diffraction fluorescent beads that serve as fiduciary marks to measure gel deformation. In addition, i^3^Neurons were labelled with a membrane dye to enable segmentation and tracking of cell features (Fig. 7c, d). High resolution imaging and force field reconstruction using the fast boundary element method (FastBEM)^41^, which suppresses noise without underestimating traction magnitude, revealed that traction stresses were localized near the edge of i^3^Neuron growth cones as well as along filopodia. These traction stresses were frequently transient (Fig. 7d, e) and in i^3^Neurons localized peak traction stress values generally remained below 100 Pa.

In DCX-mEmerald i^3^Neurons, we observed that local increases in traction stress at the growth cone edge coincided with bending of DCX-coated MTs (Fig. 7c; Video 6). Because this suggested that these DCX-coated growth cone MTs may provide mechanical resistance to actomyosin-mediated pulling forces on the extracellular matrix, we next asked how i^3^Neuron-generated traction stress changed in the absence of DCX. Because the traction stress field surrounds the growth cone edge, we quantified traction stress inside a mask that extended 1.2 µm around the growth cone defined by fluorescence threshold of the membrane label. Indeed, the average traction stress in DCX -/Y i^3^Neurons was significantly reduced, and the difference in strain energy, i.e. the work growth cones exert to deform the substrate, was even more pronounced (Control: 0.29 +/- 0.19 fJ; DCX -/Y: 0.1 +/- 0.05 fJ; Fig. 7d, f; Video 7). In contrast, we did not find a significant difference in either the duration or magnitude of transient traction stresses exerted by individual filopodia (Peak force: Control: 0.2+/- 0.16 nN; DCX - /Y: 0.15 +/- 0.11 nN; Fig. 7 g) indicating that while DCX is required to stabilize the overall pulling force exerted by the entire growth cone, a loss of DCX does not directly affect localized filopodia force production.

## Discussion

DCX loss-of-function mutations account for a sizeable portion of neurodevelopmental disorders. Yet, the molecular function of DCX in human neuron morphogenesis remains unclear. By using genome editing of hiPSC-derived cortical neurons to either label the endogenously expressed DCX protein with mEmerald or abolish DCX expression altogether, we define a new functional role for DCX in growth cone advance in a physiological soft environment. Although initial studies localized DCX to different parts of neuronal cells by immunofluorescence or overexpression^42^, we now demonstrate that at endogenous expression levels, DCX is nearly exclusively associated with straight MTs in the growth cones of developing human cortical neurons, consistent with previous transient expression data in rat hippocampal neurons^43^. This polarization of DCX-MT association is highly dynamic and coincides with the appearance of F-actin rich protrusions and nascent growth cones at very early stages of neuron morphogenesis. To our knowledge, no other neuronal MT-associated protein shows such an exclusive localization to growth cone MTs. Although it is likely that the control of DCX-MT association involves DCX phosphorylation, how this gradient of DCX-MT affinity is generated remains mysterious. The C-terminal unstructured region of DCX contains an array of cdk5 sites linked to decreased MT association in vitro^25^. Although several of these sites are phosphorylated in developing cortical neurons, direct analysis of MT-binding in DCX variants in which all putative cdk5 sites were replaced with phosphomimetic residues showed only a moderate MT-binding reduction likely due to electrostatic effects that cannot explain the >10-fold gradient observed in neurons. In our earlier experiments with non-neuronal cells, DCX also remained on MTs in mitotic cells in which high phosphorylation of cyclin-dependent kinase consensus sites would be expected^6^. In contrast, a phosphomimetic mutation of S47 that in vitro is phosphorylated by PKA or MARK^26^ reduces DCX-MT binding to a much larger extent. Thus, DCX phosphorylation in the neurite, for example by MARK kinase^26,44^, or dephosphorylation in the growth cone^45^, or both could restrict DCX association to growth cone MTs. However, this is at odds with previous biosensor-based data that MARK kinase activity is highest in the growth cone^46^ and a DCX S47 phosphorylation gradient has yet to be demonstrated in human developing neurons.

Regardless of the mechanism by which DCX-MT association is controlled in developing neurons, expression of DCX-mEmerald from the genomic locus at near endogenous levels in i^3^Neurons allowed us to estimate the absolute number of DCX molecules bound to growth cone MTs. Since DCX domains do not bind the MT seam^13^, there are approximately 1500 DCX domain binding sites per µm MT. Given that each DCX molecule has two DCX domains that are both required for MT-binding^6,7^, 750 is the upper bound of how many DCX molecules can bind to 1 µm MT. We estimated that approximately 120 DCX molecules per µm are bound to growth cone MTs, which translates to DCX-mEmerald occupying 16 +/- 4% of possible binding sites on straight growth cone MTs. However, because local DCX densities might be higher and not all potential DCX domain binding sites may be available due to other geometric constraints, this estimate likely represents a lower bound of DCX binding site occupation on growth cone MTs.

By tracking lysosome movement, we find that the growth cone is a bottleneck for MT-based transport. Lysosomes are a major kinesin-1 (KIF 5) cargo in neurons and, indeed, in vitro MT-bound DCX inhibits kinesin-1 association with MTs^15^. However, in these in vitro experiments it is not known how many DCX molecules are bound to MTs although the saturation is likely high. Because most fast MT-based organelle and vesicle transport occurs along the neurite shaft with fewer than 13 DCX molecules per µm MT (i.e. <2% of binding sites occupied), it is difficult to envision how DCX can play a direct role in regulating long range transport in neurites^47,48^. For the same reason, reported effects of DCX loss on Golgi positioning are likely indirect^49,50^. However, because around 20% saturation of available DCX-mEmerald binding sites on intracellular MTs significantly inhibited outward lysosome movement in transfected RPE cells, DCX may participate in restricting intracellular transport into the growth cone. However, because loss of DCX function also resulted in shorter growth cone MTs and an increased frequency of MT depolymerization, it is difficult to distinguish direct inhibition of KIF5-mediated transport by MT-bound DCX from effects on the growth cone network of MT tracks. Nevertheless, our results indicate that as little as one DCX molecule per layer of tubulin dimers is sufficient to stabilize and protect growth cone MTs from rapid depolymerization events. In addition, because there are generally fewer MTs in the growth cone compared to 40-50 MTs we estimate in the average i^3^Neuron neurite MT bundle, DCX-mediated MT nucleation, which occurs in vitro^7^, is likely not a physiological DCX function in growth cones.

Interestingly, we found that DCX-mediated stabilization of the growth cone MT cytoskeleton had little effect on neurite outgrowth on regular stiff tissue culture substrates, but DCX -/Y growth cone advance was inhibited on soft substrates that closely mimic the stiffness of the developing central nervous system, one of the softest materials in the body^32,51^, highlighting the importance of assessing growth cone dynamics in physiological microenvironments. The classic mode of adherent cell migration proceeds through actomyosin-produced traction forces transmitted to the extracellular matrix by transmembrane, integrin-mediated adhesion sites^52^, a model that has also been adopted for how growth cones advance^53,54^. Using high resolution traction force microscopy, we find that human cortical i^3^Neuron growth cones produce average traction stresses of approximately 10-20 Pa with around ten times larger transient peak values near the growth cone edge and along filopodia. Although it is difficult to quantify absolute traction stress values and compare between different experimental studies, our measurements are similar to traction stress magnitudes exerted by rat central nervous system^55,56^ or chick spinal neurons^57^. In contrast, traction stresses observed in fibroblasts or other types of adherent cells are frequently orders of magnitude larger than what we observe in i^3^Neuron growth cones^41,58–60^. Although traction stress dynamics correlated with growth cone dynamics, the small traction stress magnitudes generated by i^3^Neuron growth cones suggests that extracellular pulling on a highly compliant material such as the soft developing central nervous system may not be sufficient by itself for persistent and effective growth cone advance.

Due to their tubular geometry, MTs are the stiffest cytoskeleton polymer and especially when crosslinked to other cytoskeleton components individual intracellular MTs can bear large compressive loads exceeding 100 pN^36^, which is in a similar range to the transient peak traction forces produced by i^3^Neuron growth cone filopodia or at the growth cone edge. We therefore propose that the DCX-stabilized growth cone MT cytoskeleton serves as an essential intracellular mechanical component to resist actomyosin-mediated contractility and stabilize growth cone advance in the absence of a stiff extracellular scaffold. Indeed, i^3^Neuron growth cones lacking DCX display an increased retrograde F-actin flow and an even further decreased average traction stress indicating a loss of coupling of pulling forces on the extracellular matrix and forward movement. The DCX-mediated stabilization of growth cone MT length and number may by itself account for a greater compression resistance of the growth cone cytoskeleton although DCX-coated MTs appear straighter and are thus likely stiffer in cells and neurons overexpressing DCX^6,43^. In i^3^Neurons, however, growth cone MT shapes are highly variable and dynamic, and we were unable to detect a clear difference in MT curvature with or without DCX. It therefore remains unclear if DCX alters the mechanical properties of growth cone MTs at physiological expression levels although an increased frequency of MT wall defects in the growth cones of neurons from DCX knockout mice indicates a DCX function beyond only protecting MTs from depolymerization^61^. DCX has been proposed to directly interact with F-actin^62,63^. While such a crosslinking activity is consistent with our results, in DCX-mEmerald i^3^Neurons we did not observe any evidence of F-actin localization and other MT-associated proteins likely crosslink the growth cone MT and F-actin cytoskeletons^64^.

Mutations in DCX or Lis1, an activating co-factor of cytoplasmic dynein^65^, are the most frequent cause of lissencephaly-spectrum neurodevelopmental disorders^3^. While both are linked to a failure of cortical neuronal migration, DCX and Lis1 mutations show an opposite anterior-to-posterior severity gradient of associated brain malformations. We now demonstrate that the underlying mechanisms are very different, and it will be interesting to see if clinical differences can be explained by the mechanistic differences in DCX and Lis1 function. While Lis1 is required for interkinetic nuclear movement and the associated migration of immature neuron cell bodies^66^, the function of DCX is restricted to the opposite end of developing neurons promoting neurite elongation and guidance by stabilizing the MT cytoskeleton that contributes to growth cone biomechanics.

## Materials and Methods

### Molecular cloning

DCX missense mutations were produced by QuickChange II (Agilent) site directed mutagenesis of a wildtype DCX-EGFP plasmid. Plasmids in which multiple cdk5 phosphorylation sites in the DCX C-terminus were replaced with non-phosphorylatable or phosphomimetic residues were cloned by Gibson assembly (New England Biolabs) of a synthetic DNA fragment carrying these changes (synthesized by Twist Biosciences) into the DCX-EGFP plasmid cut with EcoRV and SalI.

I3 nanocage constructs tagged either with a single or a tandem repeat mEmerald at the N-terminus were constructed by Gibson assembly with PCR products encoding mEmerald and a mammalian codon optimized I3-01 K129A sequence from a plasmid obtained from M. Akamatsu^23^ into a pEGFP-N1 backbone. Of note, we tested different mEmerald-tagged I3 nanocage constructs and found that attaching mEmerald to the I3 C-terminus abolished nanocage assembly. All constructs were verified by whole plasmid sequencing (Primordium Labs).

### i^3^N culture, differentiation and genome editing

i^3^N hiPSCs were cultured and differentiated into cortical i^3^Neurons as previously described^16^. To insert mEmerald at the C-terminus of the endogenous DCX locus, cells were co-transfected with a mixture of pSpCas9(BB)-2A-GFP plasmids (gift from Feng Zhang; Addgene plasmid #48138) containing two different sgRNA sequences (TTGGATGACTCGGACTCGCT and CGCTTGGTGATTCCATGTAA) targeting the DCX exon 7 just before the stop codon and a pUC19 HDR template. Because DCX is only expressed during neuron development and not in hiPSCs, to select DCX-mEmerald i^3^N clones, individual colonies were picked into duplicate glass-bottom plates and mEmerald-expressing clones were identified by spinning disk confocal microscopy. Out of 24 colonies, we isolated one DCX-mEmerald expressing line.

To generate the DCX -/Y i^3^N line, DCX-mEmerald i^3^N cells were transfected with a mixture of three pSpCas9(BB)-2A-GFP plasmids containing sgRNA sequences in exon 1 (AAGGTACGTTTCTACCGCAA, GCGGTAGAAACGTACCTTCT and ATGGGGACCGCTACTTCAAG). DCX -/Y i^3^N cells have a 22 base pair deletion and frame shift at position 157 (…AAGAAAGCCaagaaggtacgtttctaccgcaATGGGGAC…; lower case indicates the deleted sequence). All i^3^N genome editing was confirmed by genomic PCR and sequencing of the PCR amplicon as well as immunoblotting for DCX protein. Genomic DNA was isolated using a Purelink Genomic DNA Mini Kit (ThermoFisher).

### Antibodies

Mouse monoclonal anti-DCX: Santa Cruz Biotechnology Cat# sc-271390, RRID:AB_10610966; Rabbit polyclonal anti-GFP: Invitrogen Cat# A-11122, RRID:AB_221569; Mouse monoclonal anti-MAPT(tau) E-4, sc-515539; Santa Cruz Biotechnology, no RRID available^67^. For immunoprecipitation, DCX-mEmerald i^3^Neurons were lysed in 50 mM Tris-HCl pH 7.5, 100 mM NaCl, 10% glycerol and 0.5% Triton-X100 with protease and phosphatase inhibitors. The lysate was cleared by centrifugation at 14000 rpm, incubated with anti-GFP loaded Affi-Prep Protein A Resin (Biorad), and after washing DCX-mEmerald protein was eluted in SDS sample buffer. The DCX-mEmerald band at around 70 kDa was excised from the gel and phosphorylated residues were determined by mass spectrometry (MS Bioworks LLC).

### Live microscopy and image analysis

In general, i^3^Neuron live microscopy was performed 1-3 days after replating on laminin-coated glass-bottom dishes (Mattek) essentially as described^16^. SPY555 tubulin and SPY650-FastAct (Spyrochrome) were added to i^3^Neurons at a 1:2000 and 1:3000 dilution, respectively, at least 30 min before imaging, and cells were discarded after a maximum of 3 hours. MT polymerization dynamics and F-actin retrograde flow was quantified as described^16^. Similarly, RPE cells were plated on glass-bottom and transiently transfected using Lipofectamine 2000 (Thermo Fisher) or jetOPTIMUS (Sartorius).

All live microscopy was performed either with a Yokogawa CSU-X1 spinning disk confocal essentially as described ^68^ or, for most i^3^Neuron microscopy, with a CFI Apochromat TIRF 60X NA 1.49 objective (Nikon) on a Yokogawa CSU-W1/SoRa spinning disk confocal system, and images acquired with an ORCA Fusion BT sCMOS camera (Hamamatsu) controlled through NIS Elements v5.3 software (Nikon). For high-resolution imaging of dim signal, SoRa mode was combined with 2x2 camera binning resulting in an image pixel size of 54 nm.

To determine the relative amount of MT-bound DCX in transfected RPE cells, intensity profiles perpendicular to DCX-coated MTs were fitted with a 1D Gaussian function in Matlab and the MT-bound fluorescence estimated as the integral over two standard deviations from the mean (µ +/- 2σ). The amount of DCX in the same volume of cytoplasm was estimated as 4σ(offset_Gaussian_-offset_Camera_) as indicated in Fig. S1a. Lastly, the ratio of MT-bound to cytoplasmic DCX signal for all mutated DCX constructs was normalized to the wild-type DCX MT-to-cytoplasm ratio.

To determine the absolute number of DCX molecules on growth cone MTs, we first constructed a calibration curve from RPE cells transiently transfected with I3 nanocage constructs labelled with either 60 or 120 mEmerald molecules. 2D Gaussian functions were fitted to individual nanocage dots using an interactive Matlab code and the nanocage intensity measured in a 2σ radius circle around the center of the fit minus the offset of the Gaussian function in the same area (Fig. S2a). Similarly, the DCX-mEmerald intensity was measured in 14-pixel wide rectangles along growth cone MTs, corrected for local background (Fig. S2b), and the number of DCX-mEmerald molecules was calculated using the nanocage calibration curve. Importantly, mEmerald tagged nanocage and DCX-mEmerald i^3^Neuron images were acquired using CSU-W1/SoRa super-resolution spinning disk microscopy (240x magnification; 27 nm image pixel size) with identical exposure settings during the same imaging session to minimize day-to-day instrument variability.

Lysosome velocity in i^3^Neurons was tracked with interactive custom Matlab code to select and fit the centroid of lysosomes with a 2D Gaussian function in time-lapse sequences. To quantify lysosome distribution in circular RPE cells plated on fibronectin ring micropatterns containing rhodamine fibronectin (FNR01; Cytoskeleton Inc.) that were generated as described^31^, SiR lysosome fluorescence intensity was measured in concentric 5-µm rings defined by an initial threshold of the fibronectin micropattern. MT-bound and cytoplasmic DCX-EGFP signal was determine by 1D Gaussian fitting of intensity profiles across MTs as described above.

### Chemotaxis assay

Molds to cast polydimethylsiloxane (PDMS) chemotaxis chambers were generated by photolithography on an Alveole PRIMO UV-micropatterning system^31^ in two exposure steps. First, a thin layer (3-5 µm) of SU-8 2005 (Kayaku Advanced Materials) was spin-coated onto a dehydrated silicon wafer and the 10 µm microchannels were exposed with a 20x objective and a radiant exposure of 6 mJ/mm^2^ with 20% laser power and no stitching in the PRIMO Leonardo software. After post-exposure bake, a 120 µm SU-8 2100 layer was spin-coated on top of the first layer. Using 540 nm reflected light, the exposed microchannels were aligned to the mask of the media and cell channels and exposed using a 4x objective with 3 mJ/mm^2^ radiant exposure and 100% laser power. Pre- and post-exposure bakes were according to the manufacturer’s specifications. The exposed SU-8 wafer was developed in 1-methoxy-2-propanol acetate (PGMEA) following standard protocols, and rendered hydrophobic by overnight vapor deposition of trichloro(1H,1H,2H,2H-perfluorooctyl) silane. Chemotaxis chambers were cut from Silgard 184 PDMS casts directly onto the SU-8 wafer or plastic replicas from the first, silane-contaminated PDMS cast. To remove unpolymerized PDMS, chemotaxis chambers were washed overnight in isopropanol and multiple dimes in ddH_2_O. Dried, plasma-cleaned^31^ PDMS tiles were adhered to plasma-cleaned #1.5 coverslips and the assembly baked for 30 min at 100°C, washed in ethanol, dried, UV-sterilized and filled with 50 µg/ml poly-D-lysine in PBS, incubated for 30 min at 37°C, flushed three times with PBS and then filled and incubated with 50 µg/ml mouse laminin PBS for 30 min, and finally flushed with maturation medium^16^. Pre-differentiating i^3^Ns were seeded in the central channel at a concentration of 10^6^ cells in 100 µl maturation medium without BDNF, and the medium in one of the side channels was replaced with complete maturation medium containing 10 ng/ml BDNF and 25 µg/ml fluorescent dextran. Stitched images of i^3^Neurons were acquired 2-4 days after seeding in both phase contrast (to count neurites that have passed through the microchannels) and fluorescence (to visualize the gradient).

### Traction force microscopy

For traction force microscopy (TFM), 400 Pa polyacrylamide gels were cast in 20 mm glass-bottom dishes (Mattek) essentially following a recent, detailed protocol^69^. Briefly, plasma-cleaned glass-bottom dishes were incubated with 40 µl 0.5% (3-aminopropyl) triethoxysilane for 30 min with agitation, washed three times with ddH_2_O, incubated with 80 µl 0.5% glutaraldehyde, washed three times with ddH_2_O and dried. 400 Pa polyacrylamide gels were prepared by mixing 147.6 µl ddH_2_O, 10 µl 20x PBS, 15 µl 40% acrylamide, 5 µl 2% bis-acrylamide, 2.4 µl far red fluorescent 200 nm polystyrene beads (Bangs Laboratories FSFR002), 20 µl fresh 1% ammonium persulfate and 0.2 µl tetramethyl ethylenediamine. 10 µl gel solution was added per glass-bottom dish and the solution covered with an 18 mm round plasma-cleaned and trichloro(1H,1H,2H,2H-perfluorooctyl) silane-treated coverglass. The glass-bottom dishes were flipped upside down to allow the beads to sediment toward the top of the gel and polymerized for 1 hour in a humidified chamber in the dark. After polymerization the dishes were incubated for at least 10 min in ddH_2_O to facilitate removal of the silanized coverglass. The gel surface was then activated by incubating with 1 ml 2 mg/ml 3,4- dihydroxy-L-phenylalanine (L-DOPA) dissolved in 10 mM Tris-HCl pH 10 for at least 30 min^70^, washed once with 10 mM Tris-HCl pH 10 and three times with PBS. The gels were coated with 200 µl of 100 µg/ml mouse laminin for at least 30 min at 37°C in a tissue culture incubator, washed with PBS and sterilized for 5 min with UV light in a biosafety cabinet before plating 10^5^ i^3^Neurons in complete maturation medium.

TFM time-lapse sequences were acquired on a CSU-W1/SoRa spinning disk confocal with a long working distance 40x 1.15 N.A. water immersion objective (MRD77410) in super-resolution mode resulting in an effective pixel size of 40.5 nm. Stress-free, decellularized reference images were acquired 15 min after treatment with 1% Triton X-100. TFM data were analyzed using the Fast BEM method with L1 regularization^41^ (https://github.com/DanuserLab/TFM). Because this method is computationally intensive, original 16-bit bead images were converted to 8-bit and downsampled by 50%. After subpixel alignment to the reference image, displacement fields were calculated with high-resolution bead subsampling and subpixel correlation and the following parameters: alpha value: 0.01; template size: 21 pixel; maximum displacement: 31 pixel. For each time-lapse sequence the optimal L1 regularization parameter was determined for the first frame and the same parameter used for all subsequent frames. This automatically determined L1 regularization parameter was between 10^-2^ and 10^-4^ for all datasets. To calculate the root mean square traction stress per growth cone, the growth cone CellMask image threshold was dilated by 1.2 µm. Assuming that displacement and force field vectors are largely aligned, the strain energy was estimated by the product of the magnitude of the displacement and force field maps divided by 2.

### Statistics

Details of statistical analysis including the type of test, p-values and numbers of biological replicates are provided within the relevant figures and figure legends. All statistical analysis was done in MATLAB (Mathworks, Inc.)., and graphs were produced in MATLAB and in Excel (Microsoft). In all figures, box plots show median, first and third quartile, with whiskers extending to observations within 1.5 times the interquartile range.

## Acknowledgements

We thank Li Gan for the i^3^N hiPSCs, Matthew Akamatsu for the nanocage constructs and William Dobyns for discussing DCX pathologies. This work was supported by National Institutes of Health grants R01 NS107480, S10 RR026758 and S10 OD028611 to T.W.

