## Supplementary Material for "Doublecortin reinforces microtubules to promote growth cone advance in soft environments"

### Supplementary Figures

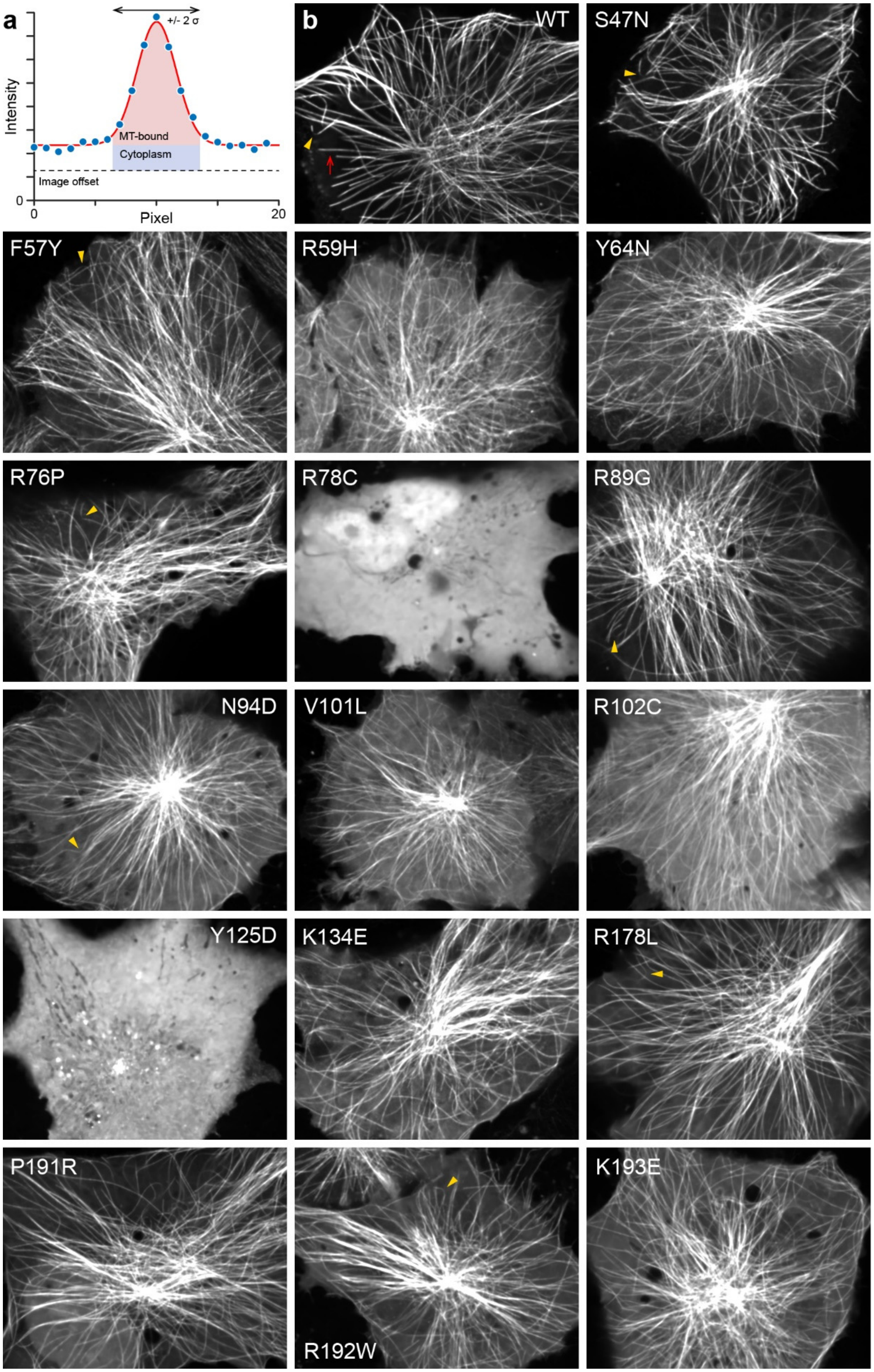

**Figure S1. Quantification of DCX-EGFP MT association. (a)** Diagram showing how MT-binding was quantified using a Gaussian fit of fluorescence intensity profiles perpendicular to the MT axis to determine the MT-bound and cytoplasmic fraction of DCX-EGFP constructs. **(b)** Representative images of all pathogenic DCX-EGFP missense mutations tested in transiently transfected RPE cells. The red arrow indicates the position of the intensity profile in **(a)**. Yellow arrows highlight MT segments with high local curvature.

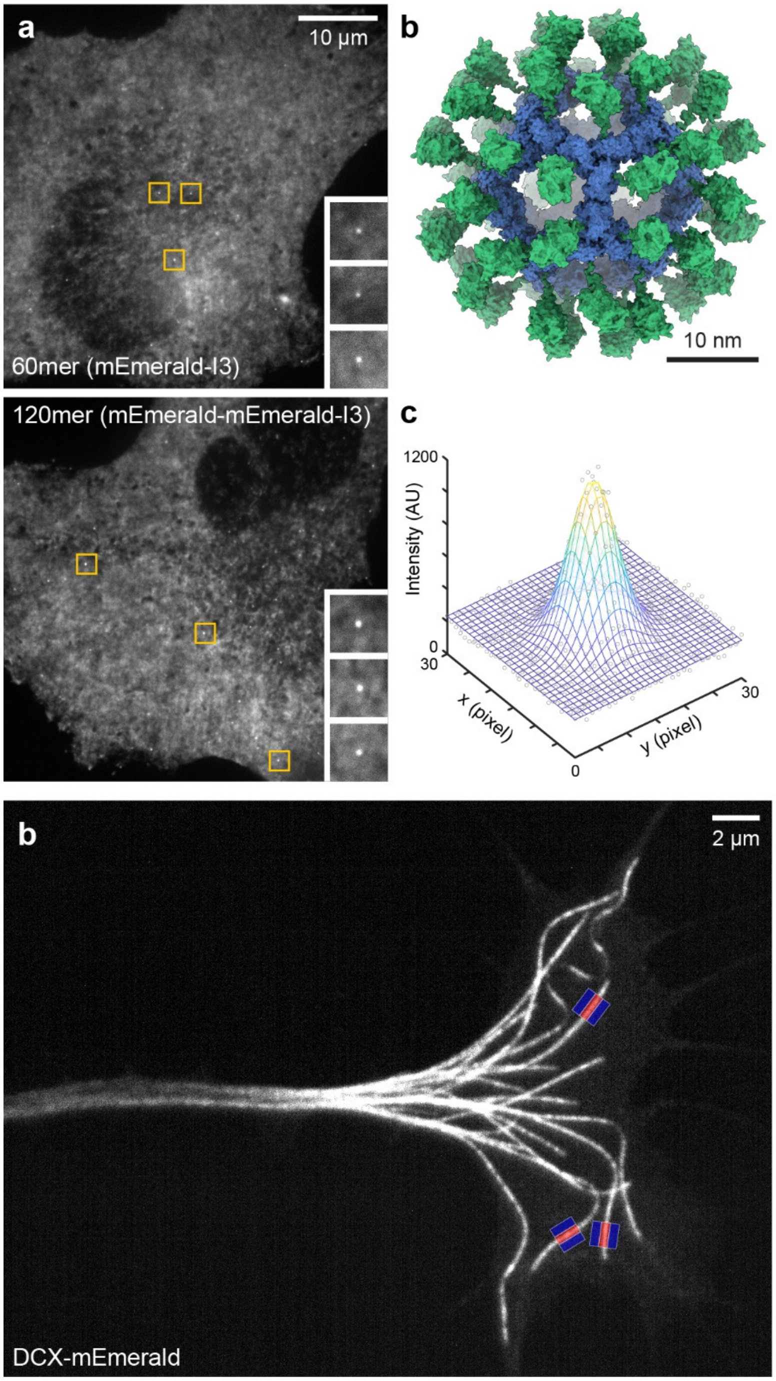


**Figure S2. Strategy for measuring the number of DCX-mEmerald molecules on growth cone MTs. (a)** Representative images of RPE cells transfected with the indicated I3 nanocage constructs carrying either 60 or 120 mEmerald molecules. Insets show the indicated regions at higher magnification. **(b)** Alphafold2 model of the mEmerald-I3 nanocage indicating that the N-terminal tag points to the outside. **(c)** Example 2D intensity profile and Gaussian fit of an individual nanocage dot. To generate the calibration curve in Fig. 3e, the integrated image intensity in a circle with a two standard deviation radius around the center of the Gaussian peak was calculated for each nanocage dot subtracting the cytoplasm background as calculated by the offset of the Gaussian fit. **(d)** Representative image of a DCX-mEmerald $i^3$Neuron growth cone acquired on the same day and with the same acquisition settings as the calibration images. Overlaid rectangles indicate regions to measure the sum of DCX-mEmerald fluorescence intensity on straight growth cone MTs (red) and local background regions (blue) on either side.

**Video legends**

**Video 1.** Time-lapse microscopy of DCX-mEmerald i$^3$Neurons labelled with SPY555-FastAct within hours after plating on a laminin substrate in very early stages of neuron morphogenesis. The total length of the recording is 8 hours and highlights the rapid incursion of DCX-mEmerald-coated MTs into nascent F-actin protrusions as well as rapid dissociation of DCX-mEmerald from MTs when a nascent growth cone retracts. Elapsed time is shown in hours:minutes:seconds.

**Video 2.** Time-lapse microscopy of DCX-mEmerald i$^3$Neurons labelled with SPY555-FastAct showing outgrowing neurites during early stages of neuron morphogenesis. The total length of the recording is 6 hours and highlights rapid DCX-mEmerald recruitment to MTs in collateral branches. The bottom panel only shows the DCX-mEmerald channel to better visualize few MTs in transient branches. Elapsed time is shown in hours:minutes:seconds.

**Video 3.** Time-lapse microscopy of DCX-mEmerald i$^3$Neurons labelled with SPY555-tubulin showing buckling of growth cone MTs and DCX-mEmerald dissociation from MT segments with high curvature. Elapsed time is shown in minutes:seconds.

**Video 4.** Time-lapse microscopy of lysosome dynamics in DCX-mEmerald i$^3$Neurons. Note that DCX-mEmerald were only captured every 10 frames to minimize photobleaching. Elapsed time is shown in minutes:seconds.

**Video 5.** Time-lapse microscopy of growth cone MT dynamics labelled with SPY555-tubulin compearing DCX-mEmerald with DCX -/Y i$^3$Neurons. For better visualization of growth cone MTs, images are shown with inverted contrast. The speckled appearance of MTs is due to sub-saturation SPY555-tubulin labelling. Elapsed time is shown in minutes:seconds.

**Video 6.** Time-lapse traction force microscopy of control and DCX -/Y i$^3$Neuron growth cones on 400 Pa laminin-coated polyacrylamide gels. The CellMask-labelled growth cone is cyan, the polyacrylamide-embedded beads are shown in grey, and the traction stress map is shown with the same 0-120 Pa pseudo-color scale as indicated in Figure 7. Elapsed time is shown in minutes.

**Video 7.** Time-lapse traction force microscopy of a DCX-mEmerald i$^3$Neuron growth cone on a 400 Pa laminin-coated polyacrylamide gel. The CellMask-labelled growth cone is cyan, DCX-mEmerald is white, the polyacrylamide-embedded beads are grey, and the traction stress map is shown with the same 0-120 Pa pseudo-color scale as indicated in Figure 7. Elapsed time is shown in minutes.
